# Local Tumor Microenvironment Niches Correlate with Survival and Immunotherapy Response in Human Glioblastoma

**DOI:** 10.64898/2026.02.23.707276

**Authors:** Florent Petitprez, Nicola C. Graham, Sheila Webb, Gillian Morrison, Lorenzo Merotto, Jessica Webb, Yuxuan Xie, Ekin Guney, William A. Weiss, Francesca Finotello, Takanori Kitamura, Steven M. Pollard, Jeffrey W. Pollard

## Abstract

**Background:** Glioblastoma (GBM) is an aggressive form of primary brain cancer. Recent efforts to characterize GBM using single-cell or spatial transcriptomics have revealed tremendous intra-tumoral heterogeneity among malignant cells and across different tumor areas. However, most studies have focused on malignant cells, and the spatial and cellular heterogeneity of the tumor microenvironment (TME) remains poorly understood. Consequently, it is unclear how TME compositions and organizations influence clinical outcomes for patients.

**Results:** Spatial transcriptomics on 25 tumors, integrated with single-cell RNA sequencing and histological analyses, enabled the estimation of cellular composition within the TME across 46,000 spots (55-μm wide). This analysis identified spatial associations between mesenchymal-like cancer cells and monocyte-derived macrophages and revealed six unique niches (N1-N6) characterized by distinct cellular composition and biological pathway activations. Deconvolution of large-scale bulk RNA-seq datasets using a spatial transcriptomics reference revealed that tumor niche composition significantly associated with patient survival and response to immunotherapy. Specifically, the N1 niche, characterized by the enrichment of mesenchymal-like cancer cells and monocyte-derived macrophages alongside hypoxic signature, was associated with lower overall survival. In contrast, the N5 niche, characterized by the enrichment of oligodendrocyte progenitor-like cancer cells and microglia-derived tumor-associated macrophages, was associated with longer patients survival. Analysis of data from patients treated with immunotherapy identified that niches N1 and N3, characterized by mixed mesenchymal-like and astrocyte-like cells, were associated with poor response to anti-PD-1 immune checkpoint inhibitors.

**Conclusions:** Our results demonstrate that GBM exhibits significant spatial heterogeneity within the TMEs, comprising distinct niche classes, and that the specific niche compositions are associated with overall survival and immunotherapy responsiveness. Our results suggest incorporation of TME niche categories as biomarkers for risk stratification and therapeutic decisions for GBM patients.

## Introduction

Glioblastomas (GBMs) are aggressive grade 4 primary brain tumors. Over 200,000 new cases are diagnosed globally every year, with 5-year survival rate below 5% [1]. Effective treatment options in GBM have remained elusive, with major challenges to both targeted therapies and immunotherapies (e.g. heterogeneity, blood-brain permeability and an immune-suppressive tumor microenvironment [2]). While immunotherapies, in particular immune checkpoint blockade (ICB), have been highly effective in many cancer types, these have generally failed in GBM [3,4]. This can be explained by the largely immunosuppressive tumor microenvironment (TME), particularly myelosuppressive cells, and challenges of ICB antibodies accession in the brain. Classical markers, such as high tumor mutational burden, specific genomic profiles, or overall immune infiltration, fail to predict subsets of ICB responders [5,6]. There is an urgent need to fully define the cellular composition and heterogeneity of tumors to support new understanding, therapeutic strategies and possibilities for patient stratification as new agents emerge.

Several efforts have been made to characterize and refine the genetic and transcriptional heterogeneity of malignant cells in GBM. In the most widely accepted molecular classification of GBM cancer cell subsets, four major cell states have been characterized: Neural progenitor cell-like (NPC-like), Oligodendrocyte Progenitor Cell-like (OPC-like), Astrocyte-like (AC-like) and Mesenchymal-like (MES-like)[7]. Varying proportions of each of these phenotypic states are present within individual tumors. MES-like has been associated with a lower survival of patients [8], as well as with a higher immune infiltration and hypoxia [7,9]. However, a major challenge is the genetic, epigenetic and cell state plasticity across these distinct differentiation and signaling states – particularly immune evasion programs [10].

Beyond the intrinsic heterogeneity of cancer cell genomes and epigenomes, TMEs are complex, highly heterogeneous and dynamic. Non-tumor cells include a wide variety of immune cells, vascular cells and other stromal cells [11,12], as well as neuronal and glia interactions at the infiltrative tumor margin. Each of these non-tumor cell types exhibits strong phenotypic inter-and intra-tumor heterogeneity that is impacted by tumor cell interactions [13]. These layers of heterogeneity are a major challenge to rational therapeutic strategies. In addition to the frequency of these cell types, their spatial organization within the tumor has also recently emerged as a crucial aspect to understanding GBM biology, initiation, progression and response to therapy [9,14–16].

The spatial organization of tumor microenvironments in GBM has not yet been fully elucidated, particularly local immunosuppressive TMEs and niches, but is essential to identify new rational therapeutic approaches. To address these questions, we integrated spatially-resolved transcriptomics with single-cell RNA sequencing (scRNA-seq), histology and multiplex immunostaining, to establish high resolution maps of cell type densities and define specific local microenvironments and niches. We stratify the GBM TME into six classes. These display distinct cellular compositions and dominant biological pathways. Deconvolution of bulk RNA-seq data allowed us to evaluate the relative abundance of these niches in large cohorts of GBM tumors and reveals an association with survival. Similar approaches on available data from patients treated with ICB revealed that some spatial niches are likely indicators of responsiveness to anti PD-1 therapies.

## Results

### Sample-wise mapping of the tumor microenvironment in GBM

To evaluate the cellular composition and spatial organization of the TMEs of GBM, spatial transcriptomics (ST) data were generated from duplicates of n=8 human IDH wild-type glioblastoma tumors using the 10x Genomics Visium technology. After quality control, 14/16 samples were deemed analyzable (UoE cohort). Additionally, we accessed data for n=11 samples from [9] (UKF cohort). **Figure 1a** describes the process to estimate cell type densities from ST data. The number of cells in each spot was first estimated using segmentation on H&E-stained images of the tumor sections. Next, spatial deconvolution approaches were used to estimate cell fractions in each spot using a publicly available single cell RNA sequencing (scRNA-seq) atlas [13]. By combining these data, fine maps of the GBM TME can be established, expressed in number of cells per spot, a measure akin to absolute cell densities since all spots are the same size.

**Figure 1:**
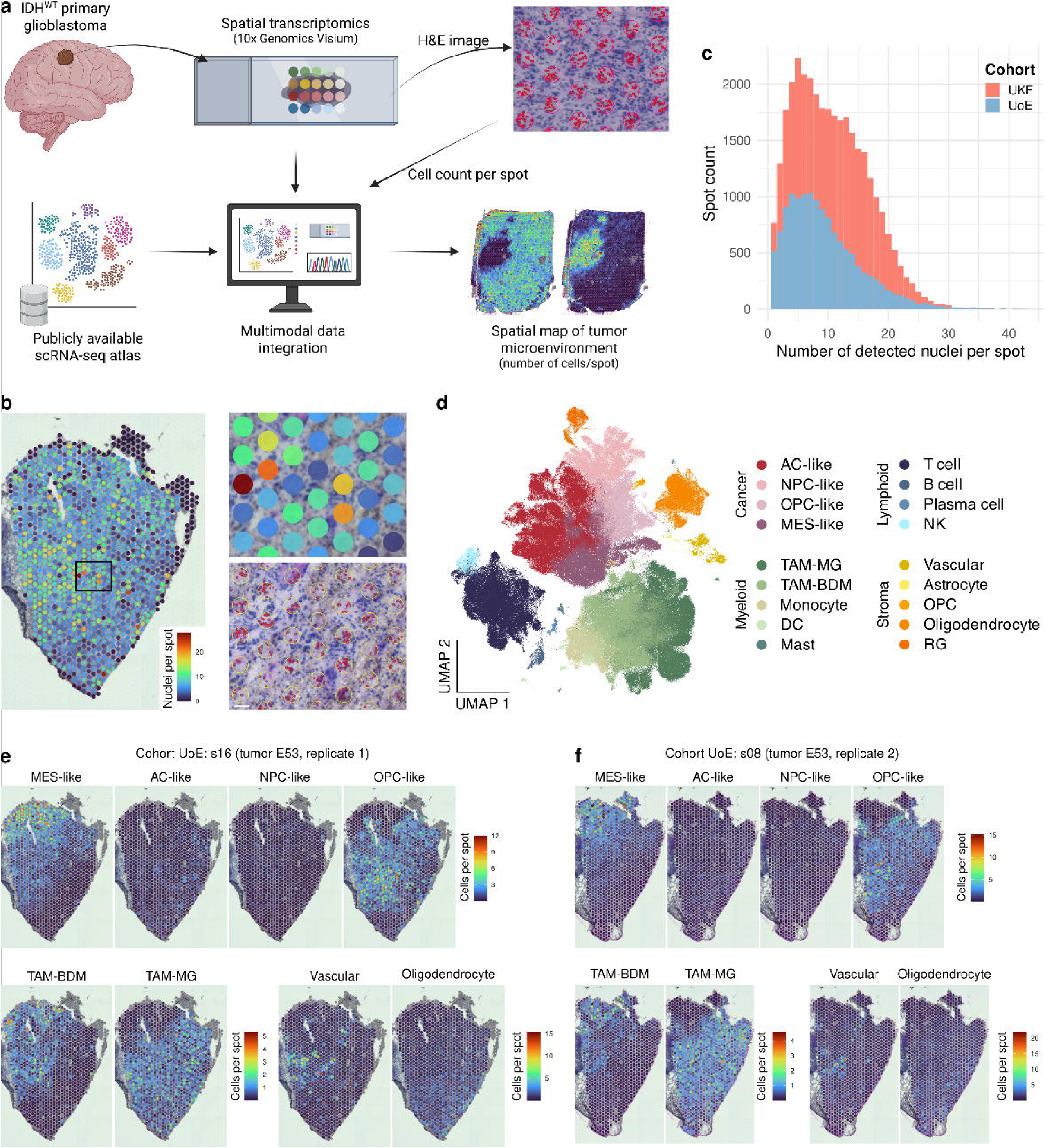
Multimodal data integration to establish TME maps. **a** Schematic representation of method to integrate spatial transcriptomics, single-cell RNA-seq and histology to draw maps of the TME expressed as estimated cell type counts per spot. **b** Representative image showing the spatial distribution of the number of detected nuclei per spot (left), Right: high magnification of an example area (top), and corresponding nuclei segmentation (bottom), with spots indicated by yellow circles and detected nuclei outlined in red. **c** Distribution of the number of detected nuclei per spot in cohorts UoE and UKF. **d** UMAP representation of the reference single-cell atlas, colored by annotated cell type. **e**, **f** Representative spatial distribution of cell types in two replicates of tumor E53 (cohort UoE). Spatial plots are colored by estimated number of cells of each cell type per spot. Color code corresponding to each plot shown on right.

To evaluate the number of cells in each spot, nucleus image segmentation and automated cellular quantification were run in areas corresponding to the spots on the image from the corresponding H&E-stained tissue section used to generate spatial transcriptomics (**Figure 1b**). The number of nuclei per spot was used as a proxy for cell count. Both cohorts exhibited similar distributions of cell counts. Median cell count was 9 cells, with a maximum of 43 cells/spot (**Figures 1c**). Plotting this cell density measure onto tissue images revealed that the heterogeneity of nuclei count per spot was also found within each sample (**Figure 1b**).

The fractions of cell types represented in each ST spot were estimated through deconvolution of each spot using Robust Cell Type Deconvolution (RCTD) [17], with the publicly available scRNA-seq atlas by Ruiz-Moreno et al. [13] as a reference, with manually curated cell type labels (**Figure 1d**). These cell types include all four types of cancer cells: AC-like, NPC-like, OPC-like and MES-like; five subsets of myeloid cells: Tumor Associated Microglia (TAM-MG), Tumor-Associated Macrophages Bone marrow-derived (TAM-BDM) [18], monocytes, dendritic cells (DC) and mast cells; four lymphocyte populations: T cell, B cell, plasma cell and NK cell; and five stromal populations: vascular (including endothelial, mural and vascular-adjacent cells), astrocyte, Oligodendrocyte Progenitor Cell (OPC), oligodendrocyte and Radial Glia (RG).

Finally, the fractions for each spot were multiplied by cell count estimates to estimate number of cells of each cell type per spot. Comparing results between duplicates revealed excellent reproducibility (**Figure 1e,f**). This was confirmed quantitively by comparing the estimates distributions with Kolomogorov-Smirnov tests. Representative comparisons for MES-like cancer cells and TAM-MG are shown in **Figure S1a-b**.

To validate the approach, a specialized pathologist (EG) labeled vessels and vascular-adjacent areas (EG, **Figure S2a-b**) and compared these with the estimated counts of vascular cells (**Figure S2c**). As expected, vascular cell density was significantly higher in vascular areas than in vascular-adjacent and further areas. This agreement between estimated cell type densities and pathologically visible structures validates the approach presented in this study.

### MES-like cancer cells are spatially segregated and associated with bone marrow-derived macrophages

An initial visual inspection of the TME maps indicates that tumors were comprised of varying proportions of the diverse GBM transcriptional subtypes and confirms the diversity of cell states [7,19]. Importantly, these states were found to be organized in specific spatial microenvironmental niches. MES-like cancer cells segregate in a different niche compared to other cancer cell subtypes, whereas AC-like and OPC-like seemed intermixed within the same spatial locations (**Figure 2a**).

**Figure 2:**
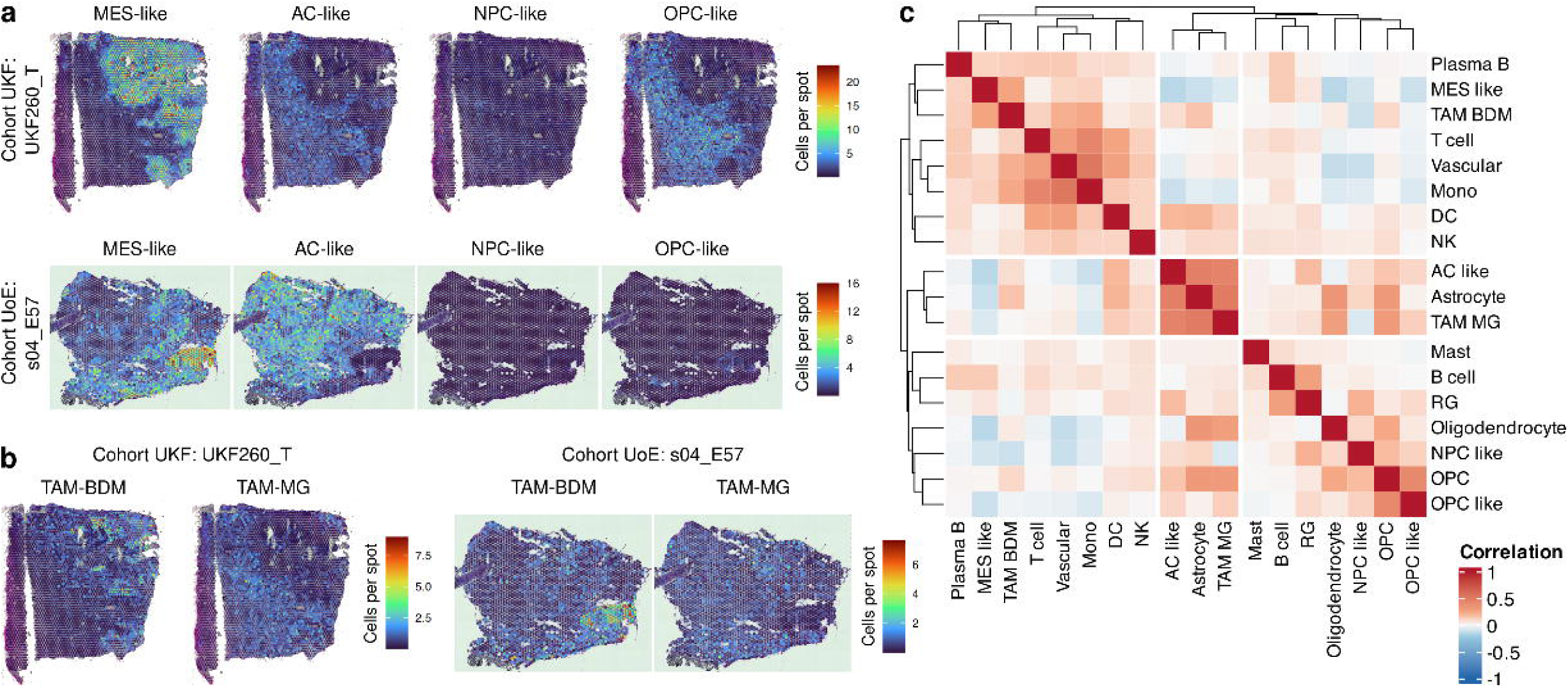
Subsets of cancer cells and different TAMs ontology occupy distinct niches. **a** Representative images showing spatial distribution of cell counts per spot for cancer cell subtypes in samples UKF260_T (up) and s04_E57 (bottom). **b** Representative images showing spatial distribution of cell counts per spot for TAMs ontologies in samples UKF260_T (left) and s04_E57 (right). **c** Pearson correlation matrix of cell type counts across all spots in both cohorts.

This segregation also applies to tumor-associated macrophages (TAMs), which segregate depending on origin. Newly recruited, bone marrow derived macrophages (TAM-BDM) were predominantly in MES-like areas, while TAMs derived from tissue-resident microglial cells (TAM-MG) populated AC-like and NPC-like areas of tumors (**Figure 2a-b**).

These patterns were then evaluated quantitatively by estimating the correlation between cell type densities across all spots over both cohorts (**Figure 2c**). This confirmed an anti-correlation between MES-like and other cancer cell subtypes, as well as a preferential association between MES-like and TAM-BDM, while TAM-MG tightly correlated with AC-like cancer cells. This is consistent with previous reports indicating the MES subtype is associated with TAMs, and may even be a cancer cell state induced by the local TAM [8,20]; positioning bone marrow-derived TAMs, rather than microglial-derived TAMs, as major orchestrators of the spatial microenvironmental niche.

This analysis also hinted at a potential perivascular, MES-like enriched niche with monocytes infiltration given the correlation with vascular cells (**Figure 2c**).

### Cross-sample integration establishes six categories of local TME niches

Previous analysis showed at least two clear niches: MES-like enriched and MES-like excluded. However, it is likely that more precise niches can be described. Therefore, a more in-depth analysis was conducted by combining all available samples to increase representativity and diversity, instead of analyzing each sample separately. Through cell count-weighted spot deconvolution, all samples were subjected to a common scale which bypasses the yet-to-be refined batch-effect correction in spatial transcriptomics analyses. This enables insights into cohort level cross-samples analyses of GBM niches. The data from all 25 samples were therefore integrated at the cell count estimate level, forming a total of 46,293 spots. A UMAP projection of all these spots based on the cell type count estimates showed that replicates from the same tumors tend to cluster together (**Figure S3a**), thereby suggesting that batch effect is not dominant. By projecting on the UMAP the cell counts (**Figure S3b**) and the estimates for different cancer cell subtypes (**Figure S3c**) and TAMs (**Figure S3d**), it was found that the organization of the UMAP seems to be mostly influenced by cancer cell type rather than by overall cell count.

Following this observation, spots were clustered to establish groups of local TME niches with similar cell type composition and count. Spots were clustered using k-means with k=6 on log-transformed cell count estimates. The optimal number of clusters was determined based on silhouette and gap statistic methods (**Figure S4**). The clusters, thereafter named niches, are projected unto the UMAP in **Figure 3a**.

**Figure 3:**
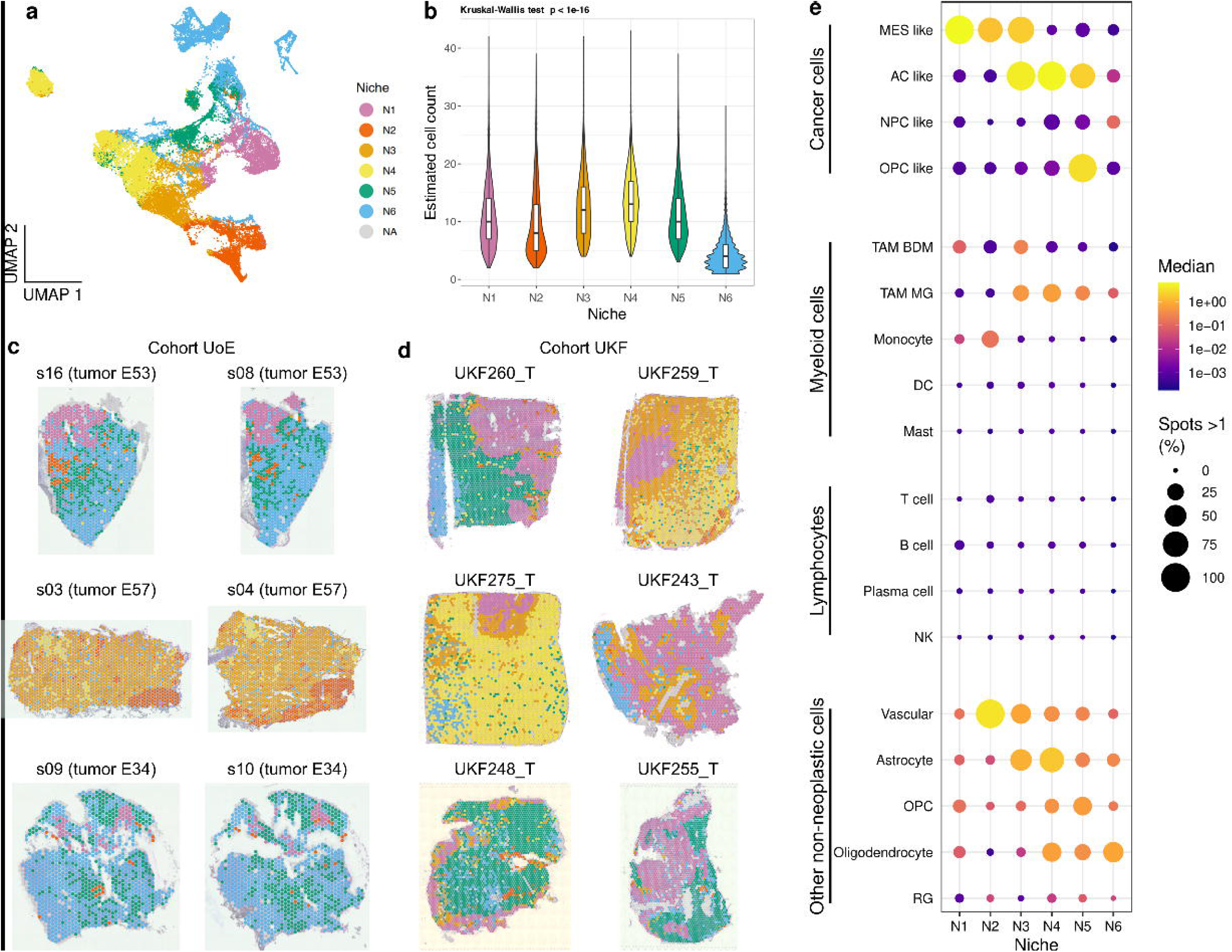
Cross-sample definition and composition of spatial TME niches. **a** UMAP representation of all spots cross both cohorts based on cell type counts estimates, colored by niches defined by k-means clustering. **b** Distribution of the number of detected nuclei per spot in all six niches. **c** Representative spatial plots representing niches spatial distribution in 3 samples from cohort UoE. Plots on left and right in each line correspond to replicates from the same tumor. **d** Representative spatial plots representing niches spatial distribution in 6 samples from cohort UKF. **e** Dotplot representing the median cell count for each cell type in each niche (color) and the percentage of spots of each niche having an estimated cell count of at least 1 (size of spots).

We first evaluated differences in overall cell density between niches (**Figure 3b**). Niches N1 to N5 seem to have broadly similar number of cells, but N6 was seen to have significantly lower number of cells. This niche likely groups spots with low overall cell counts.

By projecting the niches onto the tissue image of samples (**Figure 3c-d**), it was observed that the distribution of niches was highly organized regionally. Comparison of replicates of the same tumor (**Figure 3c**) again emphasized the replicability of the process, with extremely similar maps between replicates. This observation validates the high degree of reproducibility of the approach as well as the cross-sample integration. This also highlights that these niches do recapitulate local TME structures that are maintained over sequential tissue sections.

### Cellular composition of niches

To interpret the distinct identified niches, their cellular composition was determined on all cell types (**Figure 3e**). Focusing on the range of cancer cell states (**Figure 3e, Figure S5a**), it was observed that niches N1 and N2 predominantly harbor MES-like cancer cells and limited other cancer cell types, reminiscent of the observations of **Figure 2a-b**. Niche N3 is composed of both MES-like and AC-like cancer cells.

Niche N4 has a large predominance of AC-like cells, with, in some spots, presence of NPC-like and OPC-like cells. N5 is dominated by OPC-like cells, with the presence of some AC-like and NPC-like cells. N6, as explained above, mostly groups spots with low cellularity.

In terms of immune cells, there were limited numbers of lymphocytes (**Figure 3e**), as is usually the case in GBM. Macrophages of the two origins, microglial and bone-marrow derived, show a remarkably distinct distribution between niches (**Figure 3e, Figure S5b**). TAM-BDM are especially present in N1 and N3. In contrast, microglia-derived TAM-MG are especially abundant in niches N3, N4 and N5. This again corroborates the observations of **Figure 2a-b**; i.e., TAM-BDM are predominantly found in MES-like areas, while TAM-MG preferably locate AC-like and NPC-like areas. Niche N2 also exhibits the presence of monocytes (**Figure 3e, Figure S5c**), found only rarely in other niches. Other myeloid cell types (DC and mast cells) were only rarely identified.

Strikingly, niche N2 deviates from the profile of all other niches by displaying by far the highest density of vascular cells (**Figure 3e, Figure S5d**). Astrocytes were found to be most abundant in niches N3 and N4; OPC highest in niche N5; oligodendrocytes were predominantly found in niches N4, N5 and N6; RG were only scarcely identified in our dataset and did not exhibit strong differences between niches.

### Protein-based validation of spatial niches

To validate these observations at the protein levels, we conducted both spatial transcriptomics and immunostaining on paraffin sections from the same two tumors. Applying the aforementioned process to these spatial transcriptomics samples, we estimated cell type densities (**Figure 4a**) and subsequently assigned each spot to the nearest niche based on a centroid-based distance metric (**Figure 4b**). Additional tissue sections were stained for cancer cells and macrophage subsets using markers: SOX2, CD44, IBA1, TMEM119 and CD14. This antibody panel enables to annotate cells into MES-like cancer cell (SOX2^+^CD44^+^), non-MES like cancer cell (SOX2^+^CD44^-^), TAM-MG (IBA1^+^ or CD14^+^ and TMEM119^+^), TAM-BDM (IBA1^+^ or CD14^+^ but TMEM119^-^), or uncharacterized. Consistent with our spatial transcriptomic findings, TMEM119^+^ TAM-MG were predominantly observed in a niche N3-dominated tumor composed mainly of CD44^−^ non-MES-like cells, compared to a N1-dominated tumor with high presence of CD44^+^ MES-like cells (**Figure 4a-c**). Notably, the spatial distribution of cell types defined by immunostaining (**Figure 4d**) showed strong concordance with the niche assignments derived from spatial transcriptomics (**Figure 4b**). The same immunostaining was conducted on two more independent samples (**Figure S6a**) and cell types were assigned similarly (**Figure S6b**). Data from the four samples was combined to evaluate the distances between TAM-BDM or TAM-MG and the nearest MES-like or non-MES-like cancer cells. TAM-BDM were found to be located significantly closer to MES-like cancer cells compared to TAM-MG. In contrast, TAM-MG showed significantly shorter distance to non-MES-like cancer cells compared to TAM-BDM (**Figure 4e**). A cancer cell-centric analysis yielded consistent results (**Figure S6c**), aligning with the previous findings presented in **Figure 2** and **Figure 3**.

**Figure 4:**
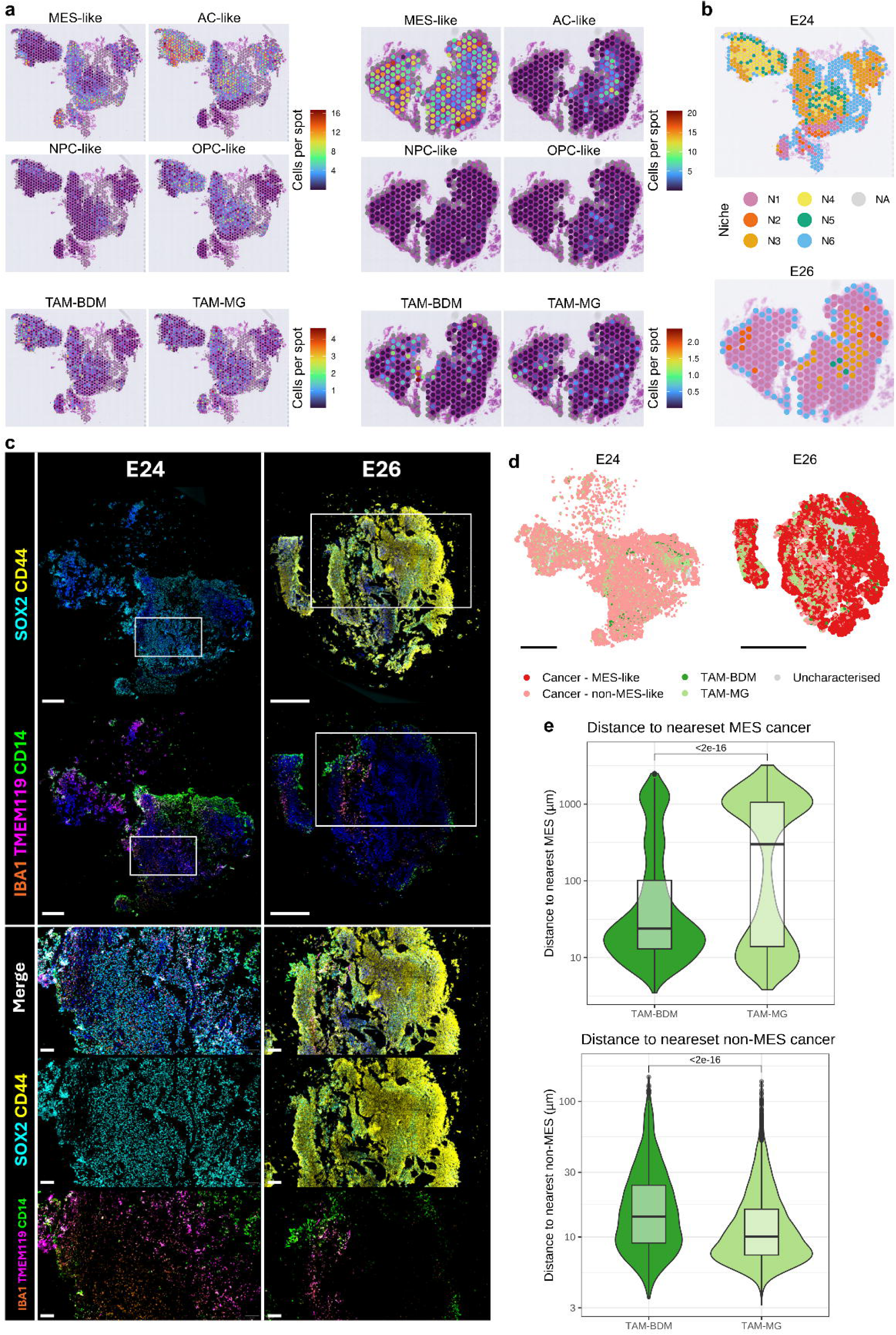
Cell-level and protein-based validation of niches. **a** Spatial transcriptomic images of two GBM samples, colored by estimated cell counts for cancer cells (top two rows) and TAMs (bottom row). **b** Spatial transcriptomics plots displaying spots colored by their assigned niches. **c** Immunostaining images of samples presented in (a), displaying the expression of SOX2 and CD44 (top) and IBA1, TMEM119 and CD14 (second line). Higher-magnification images of selected areas are also shown below. Scale bars: 500µm for whole section images, 100µm for higher magnification. **d** Map of cell types in the samples shown in (b). Individual cell type was defined by marker protein levels quantified via immunostaining. Scale bars: 1mm. **e** Distribution of distances between TAM-BDM or TAM-MG and the nearest MES-like cancer cell (left) or non-MES-like cancer cell (right). Four samples, including those presented in (b) and Figure S5 were combined. P-values are derived from Mann-Whitney tests.

### Functional annotation of niches

To determine which biological pathways were up-or downregulated in each niche, differential gene expression analysis across patient samples was performed between spots of each specific niche (N1-N6) and spots in all other niches. Significant genes were then submitted for enrichment analysis on hallmarks signatures. MES-like-enriched Niche N1 was found to upregulate notably signatures associated with hypoxia, TNFα signaling via NFκB, epithelial-mesenchymal transition (EMT), interferon γ (IFNγ) response and inflammatory response (**Figure 5a-b**) – consistent with previous reports for this signature. This niche also exhibited downregulation of oxidative phosphorylation, amongst other pathways (**Figure 5c**).

**Figure 5:**
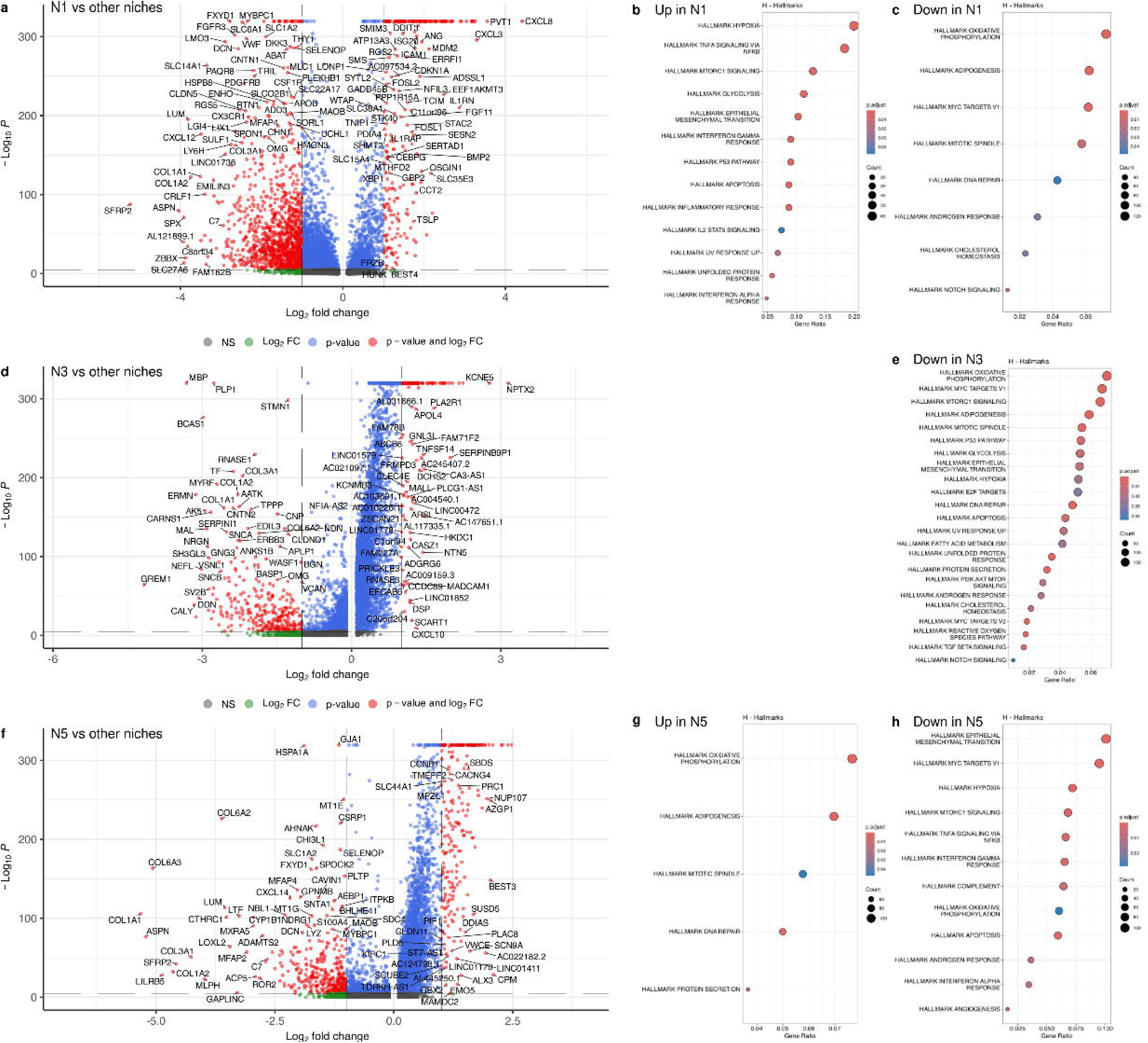
Differential gene expression and pathway enrichment in spatial niches. **a, d and f** Volcanoplot representing differentially expressed genes between niches N1 (**a**), N3 (**d**) or N5 (**f**) and all other spots. **b and g** Enrichment plots for genes identified as overexpressed in niches N1 (**b**) and N5 (**g**). **c, e and h** Enrichment plots for genes identified as underexpressed in niches N1 (**c**), N3 (**e**) and N5 (**h**).

MES-like and vascular cells-enriched Niche N2 had an overexpression of signatures related to cell cycle and proliferation processes: E2F targets, G2M checkpoint, mitotic spindle, Pi3K/AKT/MTOR (**Figure S7a-b**). It was also found to downregulate most pathways associated with niche N1, such as EMT, hypoxia, IFNγ response, TNFα via NFκB (**Figure S7c**).

Mixed MES-like and AC-like niche N3 was not found to significantly upregulate any hallmark pathway signature, but downregulate oxidative phosphorylation, several cell cycle and proliferation processes (E2F targets, mitotic spindle, Pi3K/AKT/MTOR), hypoxia, TGFβ signaling (**Figure 5d-e**).

AC-like dominated niche N4 exhibited no significantly upregulated pathway but significantly downregulated EMT, hypoxia, cell cycle processes, TGFβ signaling (**Figure S7d-e**).

OPC-like dominated niche N5 was found to upregulate among other pathways, adipogenesis, and mitotic spindle (**Figure 5f-g**), whereas EMT, MTORC1 signaling, TNFα via NFκB and IFNγ response are downregulated (**Figure 5h**).

Low cell-count niche N6 overexpressed signatures related to EMT and IFNγ, and downregulated oxidative phosphorylation, MTORC1 signaling and hypoxia (**Figure S7f-h**).

Cell-cell interaction analysis predicted that within niche N1, there could be binding between CD44 and SPP1 or FN1 (**Figure S8a**). In niche N3, this analysis predicted probable SPP1-CD44 and CD99-CD99 interaction (**Figure S8b**). In niche 5, PPIA-BSG was the interaction with the strongest probability (**Figure S8c**). Observing the expression of these genes in the scRNA-seq atlas of GBM revealed that the main potential sources of CD44 were MES-like (and potentially mast cells but these are very low in all niches), SPP1 is expressed on TAM-MG and TAM-BDM and FN1 is expressed by vascular cells (**Figure S8d**). CD99 was preferably expressed by MES-like cancer cells and, to some extent, by AC-like cancer cells, and rare T cells and NK cells. PPIA was found expressed by non-MES-like cancer cells and radial glia, while BSG was expressed by astrocytes, vascular cells and, to a lower extent, by cancer cells.

### Intra-tumor composition into spatial niches influences patient survival

Given the various cellular compositions and pathways upregulated in the different niches, it was hypothesized that the relative contribution of niches in a tumor would affect patient survival. To test this, referenced-based deconvolution of bulk RNA-seq was performed, using the present spatial transcriptomics dataset as a reference in place of the usual scRNA-seq data, using DWLS [21], accessed through omnideconv [22]. This approach was validated by simulating bulk transcriptomics by summing all counts in spots of each spatial transcriptomics data and comparing the deconvolution results with the original distribution of niches in the spatial data, considered as ground truth (**Figure S9**). This showed strong correlations between ground truth and deconvolution estimates (correlations for niches N1 to N5 ranging between R=0.84 and R=1, p-values below 2e-07), only with a lower performance to detect low cell count niche N6 (R=0.75, p=1.5e-05).

We ran this on the TCGA GBM cohort (n=151) [23]. This returned the estimated relative contribution of each spatially defined niche to the overall composition of tumors (**Figure 6a**). Integration of this data with overall survival data revealed that a higher contribution of niche N1 was significantly associated with a decreased overall survival (HR = 5.59, p = 0.019, Cox proportional hazards test) (**Figure 6b**). This association seems to be mostly related to tumors from the quartile with the highest N1 contribution (Q1-3 vs Q4, p = 0.0057, logrank test) (**Figure 6c**). While not being significant, there was a noticeable trend towards an association between niche N5 contribution and longer overall survival (HR = 0.215, p = 0.062) (**Figure 6b**). This was also related to the highest quartile of N5 contribution (Q1-3 vs Q4, p = 0.064) (**Figure 6d**).

**Figure 6:**
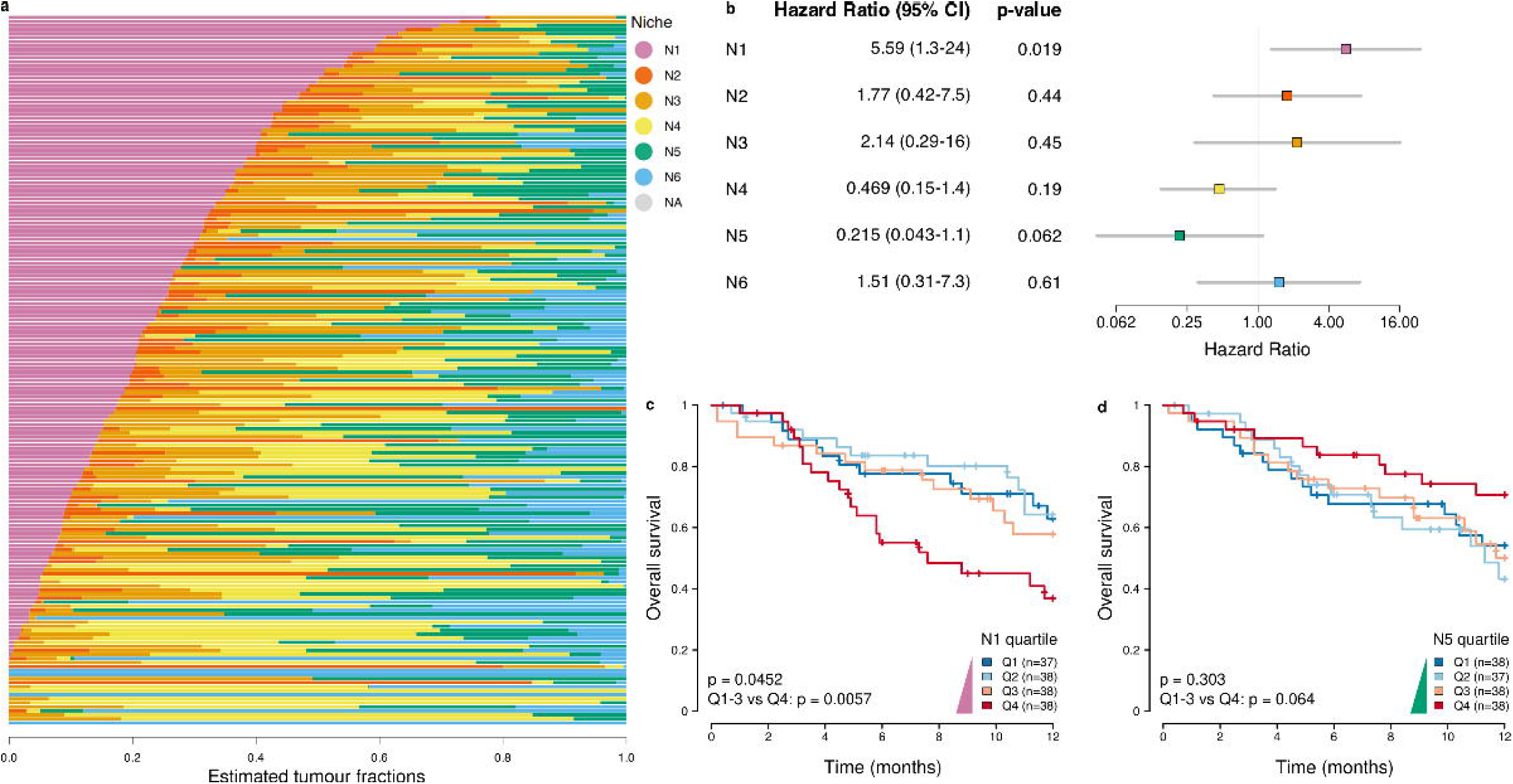
Deconvolution of bulk RNA-seq indicates prognostic relevance of niche-based tumor composition. **a** Results of niche deconvolution for all 151 tumors from the TCGA GBM cohort. Each tumor is represented by a horizontal bar and colors indicate the relative fractions of each niche. **b** Univariate Cox Proportional Hazards regression, indicating association of niches with patients’ overall survival. **c and d** Kaplan-Meier curves showing overall survival of TCGA GBM patients divided into quartiles for the contributions of niches N1 (**c**) and N5 (**d**).

### Niche contribution influences response to anti-PD-1 immunotherapy

Given the differences between niches, particularly the immune composition, it was also hypothesized that the relative abundance of each niche may influence patient response to immunotherapy. The same deconvolution approach was then applied to a dataset of patients treated with PD-1 inhibitors nivolumab or pembrolizumab [5]. This revealed that in the pre-treatment biopsy, AC-like and MES-like mixed with both TAMs ontology niche N3 was significantly more abundant in responders than in non-responders (p = 0.041) (**Figure 7a**). There was a trend towards an overrepresentation of niche N1 in responders, although it did not reach significance, likely due to the small number of samples (p = 0.093). Conversely, niche N6 was significantly more abundant in non-responders (p = 0.015). No other niche was found to differ between responders and non-responders.

**Figure 7:**
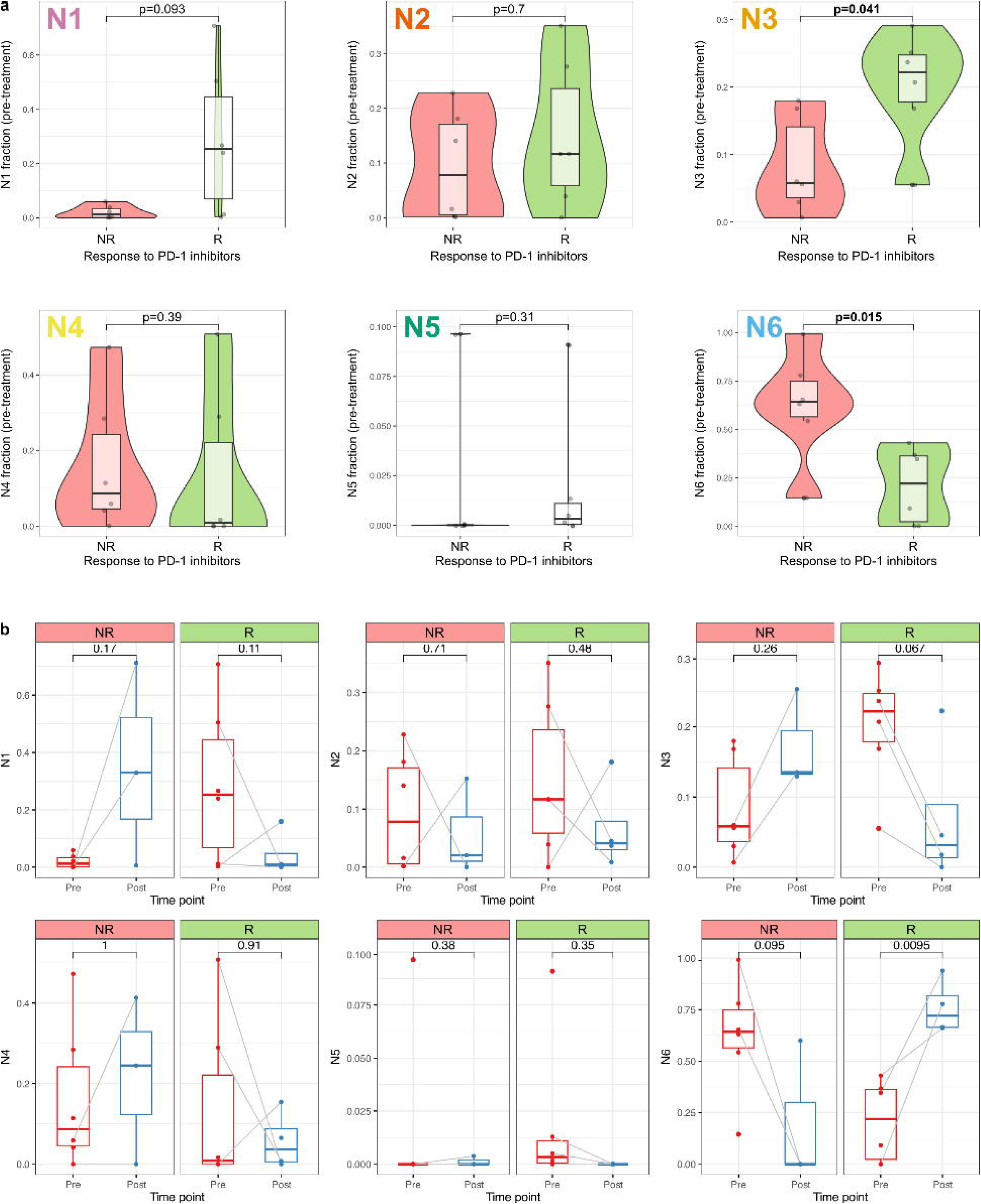
Niche composition of tumors associates with response to PD-1 inhibitors. **a** Comparison of deconvolved fractions for all niches in pre-treatment tumors between responders and non-responders. P-values are derived from Mann-Whitney tests. **b** Evolution of niches fractions between pre-and post-treatment biopsies in non-responders (No) and responders (Yes). Gray lines indicate patient-matched sample pairs. P-values are derived from Wilcoxon tests. R: Responders; NR: Non responders; Pre: pre-treatment; Post: Post-treatment.

Comparison between composition of pre-and post-treatment biopsies in responders and non-responders showed that niches N1 and N3 contribution increased during treatment in non-responders but decreased in responders, while an opposite trend was observed for niche N6 (**Figure 7b**). Overall, these data could suggest that niches N1 and N3 represent ICB-sensitive tumor regions and that, in tumors responding to the therapy, these are partially depleted, which creates an increased relative contribution of low cellularity niche N6.

In conclusion, we define six clear profiles for spatially-defined TME niches were established, with various hallmarks in terms of cancer cell type enrichment, ontogeny of TAMs, upregulated pathways, impact on survival and responsiveness to treatment with PD-1 inhibitors. These profiles are summarized in **Figure 8**.

**Figure 8:**
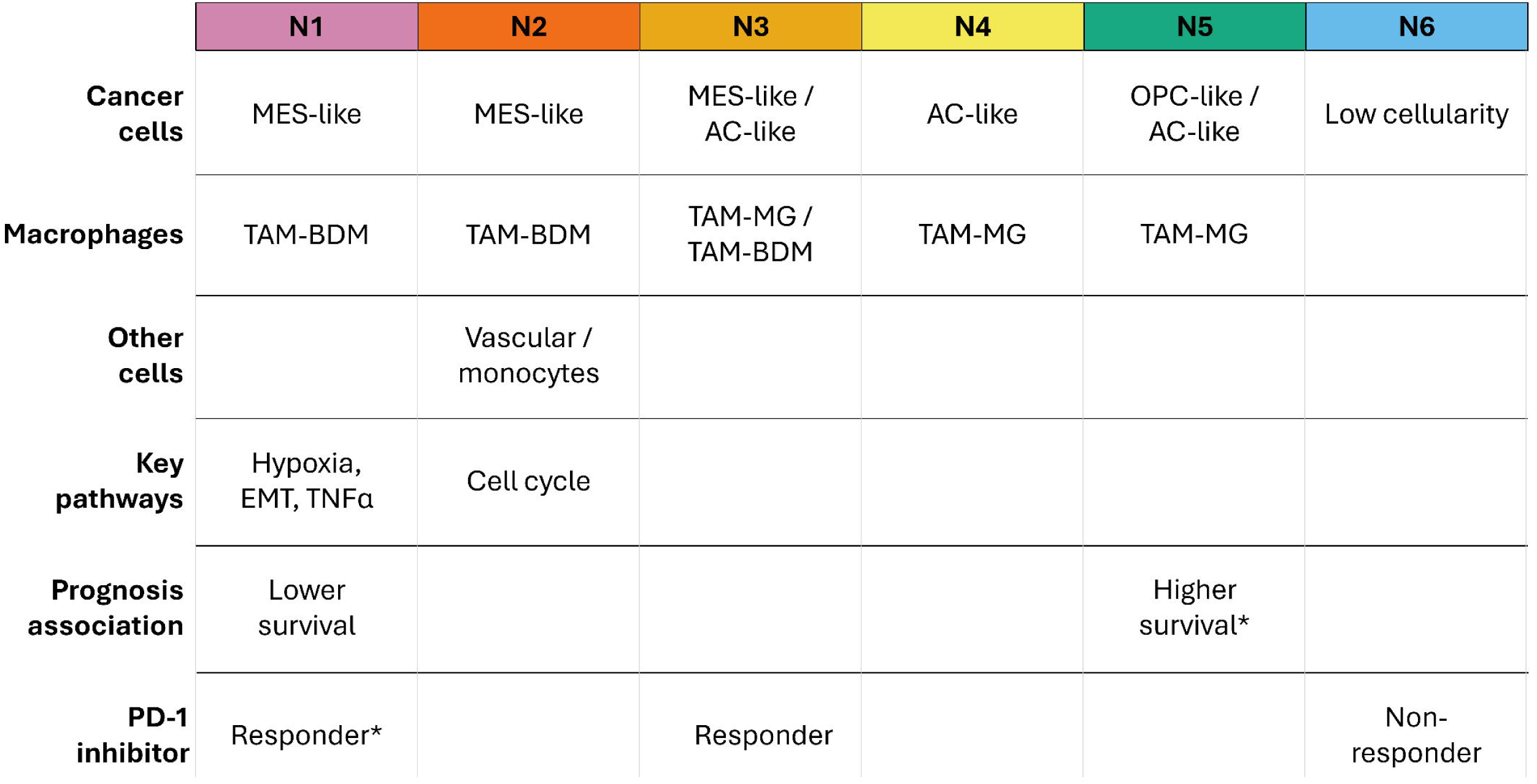
Summary of niches profiles. Elements indicated by * are derived from trends not reaching statistical significance, but with p-value below 0.1.

## Discussion

In the present study, we present an integrated approach to combine single-cell and spot-based spatial transcriptomics alongside histological image analysis, which enabled us to establish fine-grain maps of cell types densities and spatially distinct niches in GBM tumors by accounting not only for cell fractions but also for overall cell count per spot. We focused on the putatitve immune evasion niches. Using spatial transcriptomics as a reference to deconvolve bulk RNA-seq, we were able to decipher the composition of tumors into spatially-defined TME niches. This has allowed for a larger scalability of our findings by making them available from bulk transcriptomics data only. In turn, this has enabled the interrogation of clinical implications, demonstrating an association with patients’ survival and response to ICB.

### Combining spatial and single-cell transcriptomics approaches to study TMEs in GBM

Combining single-cell and spatial transcriptomics presents strong advantages compared to directly deconvoluting bulk RNA-seq with scRNA-seq references. Notably it enables the incorporation of the spatial organization of the tissue, while retaining the single-cell resolution and stronger sequencing depth offered by scRNA-seq. This enables evaluation of colocalization or spatial proximity of cell types, which is impossible from single-cell or bulk RNA-seq. Within the methods allowing this spatial deconvolution, RCTD was chosen based on methods benchmarking efforts by others [24–27].

This approach, estimating spot cellular composition from scRNA-seq reference, has already been used to provide important advances in GBM. Miller et al. used integrated scRNA-seq data and spatial transcriptomics to study immunomodulatory programs in myeloid cells [28]. Yang and colleagues used a similar approach on xenografts to determine that cancer cells exploit astrocytes ATP in the leading edge to fuel their growth [29], while Hu et al. applied this to endothelial cells subtypes [30]. However, these studies used cell fractions to analyze results.

Here, we have paired such cell fraction estimates with cell counts from the histology image to obtain cell type counts estimates, thus avoiding the potential pitfall of considering spots to be akin if they have similar cell fractions but different cell densities, which would represent quite strongly different TME areas. Others have already included histology images in their analysis, both to incorporate cell counts and to feed in deep learning approaches, thereby enabling scalability by only requiring H&E images [31]. This approach, however, is unlikely applicable when dealing with subsets of immune cells, as they are unlikely to be distinguishable from the tissue image.

### Spatial organization of the TME and segregated niches

Our approach of combining single-cell and spatial transcriptomics with histology to determine cell densities of the detailed phenotypes and subsets of immune, stromal and cancer cells, we were able to establish the spatial organization of GBM TMEs. We notably established a strong co-occurrence of bone marrow-derived TAMs with MES-like cancer cells, which seemed to occupy a niche distinct from that of other cancer cell types and TAMs deriving from brain-resident microglia. Moreover, we also found that vascularization was enriched in MES-like and TAM BDM high niche N2. These findings are consistent with previous reports involving patient-derived xenografts. Previous reports demonstrated that both origins of tumor-infiltrating macrophages (bone marrow-derived coming from the circulation and microglia originating from the adjacent tissue) have different localization patterns [32]. It was found that GBM-educated microglia-derived TAMs localized preferentially in the tumor core, whilst bone-marrow derived TAMs were found at sites of blood-brain barrier disruption.

Another study also found that macrophages and microglia exhibited compartmentalized distribution in GBM [33]. The authors notably showed that macrophages are preferentially found in areas associated with hypoxia and TNFα signaling. These are the two most upregulated pathways in our N1 niche, which exhibits a strong presence of TAM BDM. Our study adds onto this knowledge by showing that macrophages infiltrating from the circulation are mostly associated with the MES-like subset of GBM cancer cells.

In the definition of our niches, we found two separate niches enriched in MES-like cancer cells, N1 and N2, which differ by being respectively marked by hypoxia and by vasculature. Our niches also echo the work of Greenwald and colleagues [14], which also distinguished several MES-like environments. They also showed an association between macrophages and MES environments, with macrophages linked to both MES and MES-hypoxia environments. However, contrary to what we show here, they found vascular cells to be principally linked to OPC and proliferative metabolic environments, while in our work MES-like niche N2 is substantially enriched in vascular cells, and our OPC-like and OPC-rich niche N5 does not exhibit specific vascularization.

### Prognostic impact

Through deconvolution with spatial transcriptomics reference, we were able to estimate the niche composition of tumors from bulk RNA-seq. This innovative approach enables to not only evaluate the presence of each cell type, but by estimating spatially-defined niches, we can evaluate the fractions of tumors by colocalized cell types and biological pathways activated. We notably found that the MES-like and TAM-BDM rich niche N1 was significantly associated with poorer survival. This is reminiscent of the findings of Miller and colleagues [28], who showed that some tumor-infiltrating macrophages expressed a “scavenger suppressive” immunomodulatory program, associated with MES-like cancer cells. This phenotype was also associated with a lower overall survival. Zheng and colleagues [31] also showed an association between poor prognosis and hypoxia, another hallmark of niche N1. However, they also reported that poor prognosis was associated with astrocyte-like cancer cells, a feature we did not observe in our data with AC-like enriched niche N4. Instead, we found a trend towards a favorable prognostic with OPC-like and AC-like niche N5. This should be reproduced in large, independent cohorts to further ensure reproducibility

### Association with immunotherapy response

Using a similar deconvolution approach on bulk transcriptomics data from patients treated with PD-1 inhibitors, we were able to show that the fraction of low cell-count niche N6 was significantly higher in non-responders, while mixed AC-like and MES-like niche N3 was higher in tumors of responders. MES-like hypoxic niche N1 showed a trend towards a higher representation in responders. This seems in relative disagreement with the study by Miller et al., which reported a potential association between macrophages and MES-like cancer with a lower response to immunotherapy [28].

Moreover, the fractions of these niches were found to drastically change during the course of treatment, with opposite direction between responders and non-responders: niches N1 and N3 were depleted in responders, and niche N6 increased significantly in responders, while the opposite was observed in non-responders. This could suggest that parts of tumors that are classified as N1 and N3 would be responsive to immune checkpoint inhibition, being eliminated by treatment, and that the corresponding fraction could be taken over by low cellularity areas.

However, these analyses were carried out on a very limited number of patients due to unavailability of larger datasets and should be replicated in larger independent cohorts to ensure the reproducibility of our findings.

## Conclusions

In conclusion, we have evaluated the cellular composition and spatial organization of the TMEs in glioblastoma tumors. This was enabled by a robust integration of spatial transcriptomics with large single-cell RNA-seq atlas and histological image analysis. We observed a preferential infiltration of bone-marrow derived TAMs in areas enriched in MES-like cancer cells, while other areas were populated by non-MES cancer cells and had preferentially a presence of microglia-derived TAMs.

Analyzing together a large-scale dataset comprising 25 tumors and over 46,000 spatial transcriptomics spots, we defined 6 spatial niches with substantially different cellular composition and upregulated pathways (Figure 8). Niches N1 and N2 are notably composed of MES-like cancer cells and TAM-BDM, with respectively a strong hypoxic signature and a large density of vascular cells. Niche N3 has both MES-like and AC-like cancer cells, with both TAMs ontogenies. Niche N4 has AC-like cancer cells and TAM-BDM. Niche N5 has the largest OPC-like contribution, along with TAM-MG. Finally, niche N6 regroups spots with low cell density. This is one of the first reports of these niches based on finer grain annotation of both cancer cell and macrophages ontogenies, and one of the most comprehensive spatial descriptions of the TME in GBM to date.

Additionally, we established innovative deconvolution of bulk RNA-seq data using spatial transcriptomics data as a reference, instead of the single-cell RNA-seq data typically used in conventional analyses. This helped us demonstrate that niche N1 contribution is associated with significantly shorter overall survival, while niche N5 presents a trend towards longer survival. Moreover, the niche-based composition of tumors influences response to immune checkpoint inhibitors, with niches N3 and N1 being more abundant in tumors of patients benefiting from PD-1 inhibitors. These niches were subsequently found to be decreased during treatment, suggesting that they could be effectively depleted by treatment. To the best of our knowledge, this is the first report of spatially-defined areas of GBM tumors likely to differentially respond to immune checkpoint inhibitors.

The present work could pave the way to better understanding of GBM spatial organization and precision medicine by helping refine the therapeutic algorithm for each particular tumor subtype. This will subsequently permit GBM patients to receive more personalized treatment options to combat their disease.

## Materials and Methods

### Patient and tumor material

Human brain tumour samples were obtained following informed consent and all procedures received ethical approval from the NHS Health Research Authority (East of Scotland Research Ethics Service, REC reference 15/ES/0094). For cryosectioning, tissue samples were embedded in OCT and immediately stored at -80oC. Tissue was cryo-sectioned at -10°C and processed according to the recommended protocols for 10x Genomics Visium (Tissue optimization: CG000240, 10X Genomics).

### Spatial transcriptomics

#### Sample preparation, tissue imaging and library generation

Sample imaging was carried according to provider guidelines (CG000241, 10x Genomics), and permeabilization time was determined at 12 minutes. Libraries were then generated following manufacturer guidelines (CG000239, 10x Genomics) and sequenced on an Illumina NovaSeq sequencer.

#### Data processing

Manual alignment on images was performed using Loupe browser (10x Genomics) version 6.2.0. Raw read files were processed using spaceranger (10x Genomics) version spaceranger-2.0.0. The reference transcriptome was chosen as GRCh38-2020-A, and other parameters were set as default, and the manual alignments were provided. Data was then analyzed in R using package SPATA2 version 2.0.4 [34].

#### Integration with single-cell RNA-seq

Data from Ruiz-Moreno et al. [13] was accessed from CellxGene (CZI Science) at https://cellxgene.cziscience.com/collections/999f2a15-3d7e-440b-96ae-2c806799c08c, using the Core GBmap part of the atlas. Cell types provided were used as annotation_level_3. Manual curation relabeled ‘CD4/CD8’ as ‘T cell’ and both ‘Mural cell’ and ‘Endothelial’ as ‘Vascular’. Cell types having less than 100 cells were discarded from the analysis. Reference for RCTD [17] was build using the spacexr::Reference() function. Spatial transcriptomics data was read using the spacexr::read.VisiumSpatialRNA() function, and RCTD was run using mode ‘full’ to estimate fractions of each cell type.

#### Image analysis: pathologist review, nucleus segmentation, cell counts estimates

H&E images were reviewed by a trained pathologist (EG), who manually annotated spots that were considered vascular, vascular-adjacent or tumor. To determine cell counts per spot, spots were first transferred and aligned in images. Image analysis was conducted in QuPath [35] version 0.4.3. A spot alignment script was run, based on the tissue_position_list.csv and scalefactors_json.json files provided as output by spaceranger. Nuclei detection in each spot was then performed using StarDist [36] with the he_heavy_augment.pb in the StarDist QuPath extension, available at https://github.com/qupath/models/tree/main/stardist. Manual curation of the segmented nuclei was performed on all images. Spots where the nuclei detection was deemed dubious (e.g. necrotic areas, torn or folded tissue), as well as spots only partially covered by tissue were discarded from the analysis. In each remaining spot, the count of detected nuclei was used as the number cells. These cell numbers were then multiplied by the cell fractions estimated using RCTD to establish estimated number of cell of each cell type per spot.

### Clustering and definition of niches

Cell type counts estimates were log-transformed. Optimal number of clusters for k-means was determined as a consensus between the silhouette width and gap statistics methods and determined as k=6. k-means clustering was then performed on log-transformed cell type counts estimates to group spots into 6 clusters, referred to as niches.

### Differential gene expression and upregulated pathways

Analysis of differentially expressed genes was performed using Seurat [37] version 5.3.0 and the function Seurat::FindAllMarkers() with default parameters. Pathway enrichment analysis was performed separately on the list of over-and under-expressed genes for each niche using ClusterProfiler [38] and the clusterProfiler::enricher(), with the Hallmarks gene sets from MSigDb [39], version h.all.v2024.1.Hs.symbols.gmt.

### Prediction of cell-cell interactions

Cell-cell interactions in spatial transcriptomics data were performed using CellChat v2 [40]. Counts were normalized with CellChat::normalizeData(). Cell-cell interaction network was evaluated with CellChat::computeCommunProb() with distance.use set to FALSE, contact.dependent set to TRUE with contact.range to 100. Communication network was filtered with minimal spot count set to 10.

### Multiplex Immunofluorescence Staining

Multiplex immunofluorescence staining was performed on GBM FFPE sections using the Leica Bond Rx fully automated stainer platform and the OpalTM 6-plex detection system (Quanterix, NEL871001KT), according to manufacturer’s instructions. Formalin-fixed, paraffin-embedded sections (5µm) were baked at 56°C for 18 hours, deparaffinised in xylene (2 x 5minutes) and rehydrated through graded ethanol (100%, 100%, 80%, 70%, 50%) for 20 seconds each) before washed in water.

For each staining cycle, antigen retrieval was performed at 100°C for 20 minutes in either BOND epitope retrieval buffer 1 (citrate buffer pH6.0, Leica, AR0086) or BOND epitope retrieval buffer 2 (EDTA buffer, pH9.0, Leica, AR9640) according to requirements of the primary antibody. Endogenous peroxidase activity was quenched using 3% hydrogen peroxide for 10 min, followed by blocking with antibody diluent/blocking buffer (Quanterix, ARD1001A) for 30 minutes. Sections were incubated with primary antibody for 60 minutes followed by Opal Polymer HRP Ms + Rb secondary antibody (Quanterix, ARH1001A) for 30 minutes. Opal fluorophores (Quanterix) were subsequently applied at 1/150 dilution in 1x Plus Amplification Diluent (Quanterix, FP1609) for 10 min. Following each staining cycle, antibody complexes were removed by the antigen retrieval of the subsequent cycle whilst preserving the covalently deposited opal fluorophore signal.

For the final staining cycle, opal780 (FP1501) was applied using a tyramide signal amplification-digoxigenin (TSA-DIG) approach. Following secondary antibody incubation, sections were incubated with TSA-DIG (Quanterix, NEL748) for 10 minfollowed by ER1 buffer at 95°C for 20 minutes to remove the primary-secondary antibody complex whilst retaining the TSA-DIG. Sections were subsequently incubated with anti-digoxigenin antibody conjugated to opal780 (Quanterix, FP1501) at 1/25 diluted in antibody diluent/blocking buffer for 10 minutes. Following completion of all staining cycles, nuclei were counterstained with 1x Spectral DAPI (Quanterix, FP1490), and slides mounted using antifade mounting medium (Thermo, P36961).

The following panel was used: CD14 (clone SP192, Abcam, ab230903, 1/50, ER2); CD44 (clone IM7, Novus, NBP1-41266, 1/200, ER2); CD31 (C31.3+JC/70A, antibodies.com, A278329, 1/1000, ER2); SOX2 (clone EPR3131, Abcam, ab215970, 1/50, ER2); IBA1 (clone EPR6136(2), Abcam, ab178680, 1/500, ER1); TMEM119 (clone CL8714, Atlas, AMAb91528, 1/500, ER2).

### Imaging, post-processing and analysis

Whole-slide multispectral images were acquired using PhenoImager HT Vectra® Polaris™ Imaging System (Akoya Biosciences, software version 1.0.13) at 40x magnification. Phenochart (version 1.1.0) was used to generate project files and define regions on interest for batch analysis. Selected ROIs were exported to InForm (version2.7.0) where spectral unmixing was performed using the built-in spectral unmixing algorithm. The resulting unmixed component data images were subsequently stitched using Visiopharm® (version 2023.09.4.15595 x64).

Processed images were analysed using QuPath [35] (Version 0.6.0). Cells were segmented using InstanSeg [41] and phenotyped using cell classification based on threshold values of marker expression. Cell measurements were exported from QuPath and analysed using R version 4.5.1.

### Bulk RNA-seq

#### Data accession and processing

Data from the TCGA GBM [23] cohort was accessed on the cBioPortal database [42] (Glioblastoma (TCGA, Cell 2013)). Gene expression data was downloaded as RNAseq v2 RSEM and log2-transformed. Clinical data at the patient level were also accessed from cBioPortal. Bulk RNA-seq data from tumors treated with PD-1 blockade from Zhao et al. [5] were accessed from SRA with accession code PRJNA482620, as raw reads. Raw reads were aligned with STAR [43] version 2.7.8a_2021-03-08 on human transcriptome GRCh38.107. Gene counts were generated using featureCounts [44]. Data was normalized as transcripts per millions (TPM) and log2-normalized. Clinical data was accessed from the supplementary tables of [5] and correspondence between samples was obtained from the authors (personal communication).

#### Deconvolution

Deconvolution was performed using the method DWLS [21], accessed through the R package omnideconv (v1.0.0) [22]. The analysis was carried out in two steps: signature building and deconvolution. The signature matrix was built starting from the spatial counts and their niches annotation with the function “build_model_dwls”, with the parameter dwls_method = “mast_optimized”. For the signature building, spatial dataset was randomly subset to retain at most 5’000 cells per niche. Deconvolution was then performed on the TPMs and the precomputed signature with the “deconvolute_dwls” function, with the parameter dwls_submethod = “DampenedWLS”.

#### Survival analysis

The association between niches composition and survival was first assessed using Cox proportional hazards models. Survival data from the TCGA GBM cohort were censored at 12 months to measure the impact on the one-year survival. Analysis was also conducted by discretizing the niche fractions into quartile. Survival was plotted using Kaplan-Meier curves and tested with logrank tests.

### Statistical analysis

Statistical analyses were conducted with R version 4.5.1. Spatial transcriptomics analyses were conducted with packages SPATA2 version 3.1.4 and Seurat version 5.3.0. Comparisons of numerical variables between groups were conducted using Mann-Whitney tests for 2 groups and Kruskal-Wallis test for 3 or more groups, followed by Dunn post-hoc test for pairwise comparisons with Benjamini-Hochberg correction for multiple testing. Paired comparisons such as evolution of niches contribution during treatment were evaluated with Wilcoxon tests. Association between numerical variables was assessed using Pearson correlation. Comparisons of distributions were conducted with Kolmogorov-Smrirnov tests. Survival analyses were conducted using univariate Cox proportional hazards model for quantitative variables and using logrank test for categorical variables. Significance threshold for p-values was set to 0.05.

## Declarations

### Ethics approval and consent to participate

Human brain tumour samples were obtained following informed consent and all procedures received ethical approval from the NHS Health Research Authority (East of Scotland Research Ethics Service, REC reference 15/ES/0094).

### Data and code availability

The spatial transcriptomics dataset (cohort UoE) generated and analyzed during the current study is available from Figshare: www.doi.org/10.6084/m9.figshare.33605389.

All other data analyzed in this study are publicly available. Spatial transcriptomics from the UKF cohort can be accessed from Datadryad (https://doi.org/10.5061/dryad.h70rxwdmj). Single-cell RNA-seq data from the reference atlas can be accessed at https://cellxgene.cziscience.com/collections/999f2a15-3d7e-440b-96ae-2c806799c08c. Bulk RNA-seq data from the TCGA GBM cohort can be accessed from the cBio Portal (https://www.cbioportal.org/study/summary?id=gbm_tcga_pub2013). Bulk RNA-seq data from patients treated with PD1 inhibitors can be accessed from SRA (accession code: PRJNA482620).

The code used to analyse the data and generate the plots is available on GitHub: https://github.com/FPetitprez/SpatialGBM.

### Authors’ contributions

Study conceptualization and design: FP, SMP, JWP

Data acquisition: NCG, SW, GM, JW

Data analysis: FP, NCG, LM, YX, EG

Data interpretation: FP, WAW, FF, TK, SMP, JWP

Study supervision: TK, SMP, JWP

Manuscript preparation: FP, FF, TK, SMP

All authors have read and approved the manuscript.

## Acknowledgements

The authors wish to acknowledge Daniel Soong, Jo Van Ginderachter and Marianna Romano for insightful discussions in the build up to this study. Samples were processed and data generated with technical support from the IRR Histology & Multiplexing Facility, Institute for Regeneration & Repair, University of Edinburgh. The computational results presented here have been achieved in part using the resources provided by the Edinburgh Compute and Data Facility (ECDF) (http://www.ecdf.ed.ac.uk/) and the LEO HPC infrastructure of the University of Innsbruck. The results shown here are in part based upon data generated by the TCGA Research Network (https://cancergenome.nih.gov).

---

**Figure S1: Cell type numbers estimates are reproducible between replicates**

**a and b** Representative distribution of estimated counts for MES-like cancer cells (**a**) and TAM MG (**b**). Each set of 2 plots (dark and light hues for the same colors) corresponds to a pair of duplicates. P-values are derived from Kolmogorov-Smirnov tests to evaluate significant shifts in distributions.

**Figure S2: Vascular cell estimates correlate with pathologist annotation**

**a** Spatial plot of 3 representative tumors with annotations of zones as vascular, vascular-adjacent and tumor. **b** H&E details illustrating pathologist annotation of vascular (red circle), vascular-adjacent (yellow circle) and tumor (blue circle) spots. **c** Distribution of estimated number of vascular cells in each zone for each the samples presented in **a**. P-values are derived from Dunn tests with Benjamini-Hochberg correction for multiple testing. Dotted lines indicate the level of 1 cell/spot.

**Figure S3: Cross-cohort UMAP is mostly organized by cancer cell type**

**a-c** UMAP of all spots based on cellular composition, colored by total cell count (**a**), cancer types counts (**b**) and TAMs counts (**c**). Color code was restricted to the range shown to the right of each plot, and spots with higher counts were set to the max color.

**Figure S4: Determination of optimal number of niches for k-means clustering**

Using silhouette width (**a**) and gap statistic (**b**).

**Figure S5: Cellular composition of the niches**

Violinplots showing the distribution of estimated cell type counts in each of the niches for cancer cells (**a**), tumor-associated macrophages (**b**), monocytes (**c**) and vascular (**d**).

**Figure S6: Cell-level and protein-based validation of niches**

**a** Immunostaining images of GBM, displaying the expression of SOX2 and CD44 (top) and IBA1, TMEM119 and CD14 (second line). Higher-magnification images of selected areas are also shown below. Scale bars: 500µm for whole section images, 100µm for higher magnification. **b** Map of cell types in the samples shown in (b).

Individual cell type was defined by marker protein levels quantified via immunostaining. Scale bars: 1mm. **c** Distribution of distances between MES-like or Non-MES-like cancer cells and the nearest TAM-BDM (left) or TAM-MG (right). Four samples, including those presented in (a) and Figure 4 were combined. Vertical bar indicates 40µm. P-values are derived from Mann-Whitney tests.

**Figure S7: Differential gene expression and pathway enrichment in spatial niches**

**a, d and f** Volcanoplot representing differentially expressed genes between niches N2 (**a**), N4 (**d**) and N6 (**f**) and all other spots. **b and g** Enrichment plots for genes identified as overexpressed in niches N2 (**b**) and N6 (**g**). **c, e and h** Enrichment plots for genes identified as underexpressed in niches N2 (**c**), N4 (**e**) and N6 (**h**).

**Figure S8: Cell-cell interaction predictions**

**a-c** Cell-cell interaction predictions within niches N1 (**a**), N3 (**b**) and N5 (**c**). **d** Dotplot representing average expression of selected genes in cell types in scRNA-seq data.

**Figure S9: Evaluation of bulk deconvolution with spatial transcriptomics reference**

Correlation between the niche composition in spatial transcriptomics (Ground truth) and the deconvolution estimate (Deconvolution) of each niche fraction, obtained by deconvolving the pseudobulked spatial counts. Blue line and greyed area indicate linear regression and 95% confidence intervals. Dashed line indicates y=x (perfect agreement). R denotes Pearson correlation. P-values are derived from a correlation test.

